# Reduction of cybersickness during and immediately following noisy galvanic vestibular stimulation

**DOI:** 10.1101/843060

**Authors:** Séamas Weech, Travis Wall, Michael Barnett-Cowan

## Abstract

The mechanism underlying cybersickness during virtual reality (VR) exposure is still poorly understood, although research has highlighted a causal role for visual-vestibular sensory conflict. Recently established methods for reducing cybersickness include galvanic vestibular stimulation (GVS) to mimic absent vestibular cues in VR, or vibration of the vestibular organs to add noise to the sensory modality. Here, we examined if applying noise to the vestibular system using noisy-current GVS also affects sickness severity in VR. Participants were exposed to one of two VR games that were classified as either moderate or intense with respect to their nauseogenic effects. The VR content lasted for 50 minutes and was broken down into 3 blocks: 30 minutes of gameplay during exposure to either noisy GVS (±1750 μA) or sham stimulation (0 μA), and 10 minutes of gameplay before and after this block. We characterized the effects of noisy GVS in terms of post-minus-pre-exposure cybersickness scores. For the intense VR content, we found a main effect of noisy vestibular stimulation. Participants reported lower cybersickness scores during and directly after exposure to GVS. However, this difference was quickly extinguished (∼3-6 min) after further exposure to VR, indicating that sensory adaptation did not persist after stimulation was terminated. In contrast, there were no differences between the sham and GVS group for the moderate VR content. The results show the potential for reducing cybersickness with simple non-invasive sensory stimulation. We discuss the prospect that noise-induced sensory re-weighting is responsible for the observed effects, and address other possible mechanisms.

## 1. Introduction

A persistent conflict experienced in virtual reality (VR) applications is the discrepancy between multisensory cues, particularly those processed by the visual and vestibular senses. Conditions of sensory conflict are thought to generate cybersickness (CS; Hettinger et al., 1990; Keshavarz et al., 2015a,b; McCauley & Sharkey, 1992; Oman, 1990; Reason, 1978; Stanney & Hash, 1998; Weech et al., 2019; c.f. Stoffregen & Riccio, 1991), a syndrome encompassing feelings of disorientation, nausea, and discomfort (Rebenitsch & Owen, 2016; Stanney et al., 1997). Evidence acquired from neurophysiology supports a link between sensory conflict and vestibular nuclei activation (Oman & Cullen, 2014), which share connections with other emetic centres of the brain, including the nucleus tractus solitarius and parabrachial nucleus (Yates et al., 2014).

Apart from the central conflict theory, other etiological accounts of motion/cyber sickness have been investigated in detail. For instance, it has been proposed that the provocative stimulus is not sensory conflict, but the postural instability related to novel environmental conditions such as VR (Riccio & Stoffregen, 1991; Stoffregen et al., 2014; Smart Jr. et al., 2002). Additionally, the sensory conflict account has been refined by some to focus solely on mismatches between the estimated and true environmental vertical (Bles et al. 1998; Bos et al. 2008). Cohen and others (Cohen et al., 2003, 2008; Dai et al., 2010, 2007) have proposed that individual differences in the rate of decay in stored estimates of head movement (time constant of velocity storage) contribute to motion sickness. This claim is supported by evidence that procedures which modify these estimates (i.e., opposing, low-magnitude, visual-vestibular stimulation) habituates participants to nauseating conditions (Dai et al., 2011). A unifying aspect of all etiological theories is the primacy of self-motion estimation and control, which is predominately mediated by visual-vestibular integration.

Cybersickness is reduced when the absent vestibular cues during simulated self-motion are mimicked by applying skin conducted electrical current at the mastoid processes (Cevette et al., 2012; Gálvez-García et al., 2015; Reed-Jones et al., 2007). Bilateral galvanic vestibular stimulation (GVS) can produce perceived head-tilt sensations that, when delivered at the appropriate moments (e.g., navigating a sharp bend in a driving simulation), are sufficient to re-couple visual and vestibular estimates of self-motion (Fitzpatrick & Day, 2004). Here, the stimulation device and the rendering software are coupled in order to generate congruent visual and vestibular stimuli. However, galvanic stimuli can provide only a coarse replication of the complex pattern of vestibular cues that occur during physical self-motion (St George & Fitzpatrick, 2011), which reduces the practical application of the sensory re-coupling approach.

Other recent evidence has suggested that CS is counteracted by facilitating sensory re-weighting. Time-coupling a noisy (vibratory) vestibular stimulus to the occurrence of expected vestibular cues reduces sickness during VR use (Weech et al., 2018a). The addition of noise to the vestibular sense was considered to rapidly reduce vestibular cue reliability, causing visual self-motion cues to be up-weighted in return. This finding indicates that sensory re-weighting within a single experimental session can reduce CS. Vestibular noise can also enhance illusory self-motion (vection), a known correlate of CS (Keshavarz et al., 2015b; Weech et al., 2018b), likely by facilitating a similar sensory re-weighting mechanism (Weech & Troje, 2017). In the context of optimal cue integration (Carver et al., 2006; Welch & Warren, 2004; Ernst & Bülthoff, 2004; Ernst & Banks, 2002), noise applied to the vestibular system using galvanic stimulation might down-regulate the use of vestibular cues, according to the principle that the precision of a modality determines its utility (Welch & Warren, 1980; Ernst & Bülthoff, 2004). Such an outcome would reduce the impact of absent inertial cues to motion during simulated self-motion in VR. Existing evidence has confirmed a reduced reliance on vestibular cues following GVS exposure in a manner consistent with sensory re-weighting (Balter et al., 2004a, 2004b; Dilda et al., 2014; Moore et al., 2015).

The current study was motivated by the prediction that exposure to noisy vestibular stimulation applied during VR exposure will reduce CS. The proposed mechanism was via down-regulation of ‘unreliable’ vestibular cues that would otherwise signal a stationary head-in-space. Here, we exposed participants to noisy GVS while they completed VR tasks, and measured self-reported levels of CS. There were two VR tasks, classified as ‘moderate’ or ‘intense’, that differed in terms of the frequency of simulated self-motion. We took self-report measures of CS during and after exposure to VR for noisy GVS and Sham stimulation groups. We expected to observe a divergence in sickness severity for the GVS and Sham groups to emerge, consistent with sensory re-weighting. At the same time, we aimed to establish the efficacy of noisy GVS as a reductive therapy for cybersickness.

## 2. Methods

### 2.1. Participants

40 adults (23 women, average age 22 yrs ± 7.04 *SD*, range 18-56) participated in the study. We screened participants for exclusion criteria using a self-report questionnaire that was completed by individual participants prior to the study. Each participant reported normal/corrected to normal vision and reported being free of any musculoskeletal and neurological impairment, medical implants (e.g., pacemakers), vestibular or balance deficits, and that they had no history of severe motion sickness. Due to the use of rubbing alcohol to prepare the skin surface for GVS, individuals with skin allergies were also unable to take part. Participants were informed of all procedures and apparatuses and provided written consent. All experiments were performed with the approval of the University of Waterloo’s research ethics committee on human research, and were conducted in accordance with the declaration of Helsinki.

### 2.2. Apparatus and stimuli

Vestibular stimulation was provided via a galvanic vestibular stimulator (DC-Stimulator Plus, neuroConn GmbH, Germany) through 25.4 mm-diameter self-adhesive silver chloride electrodes (Figure 1a).

**Fig. 1.**
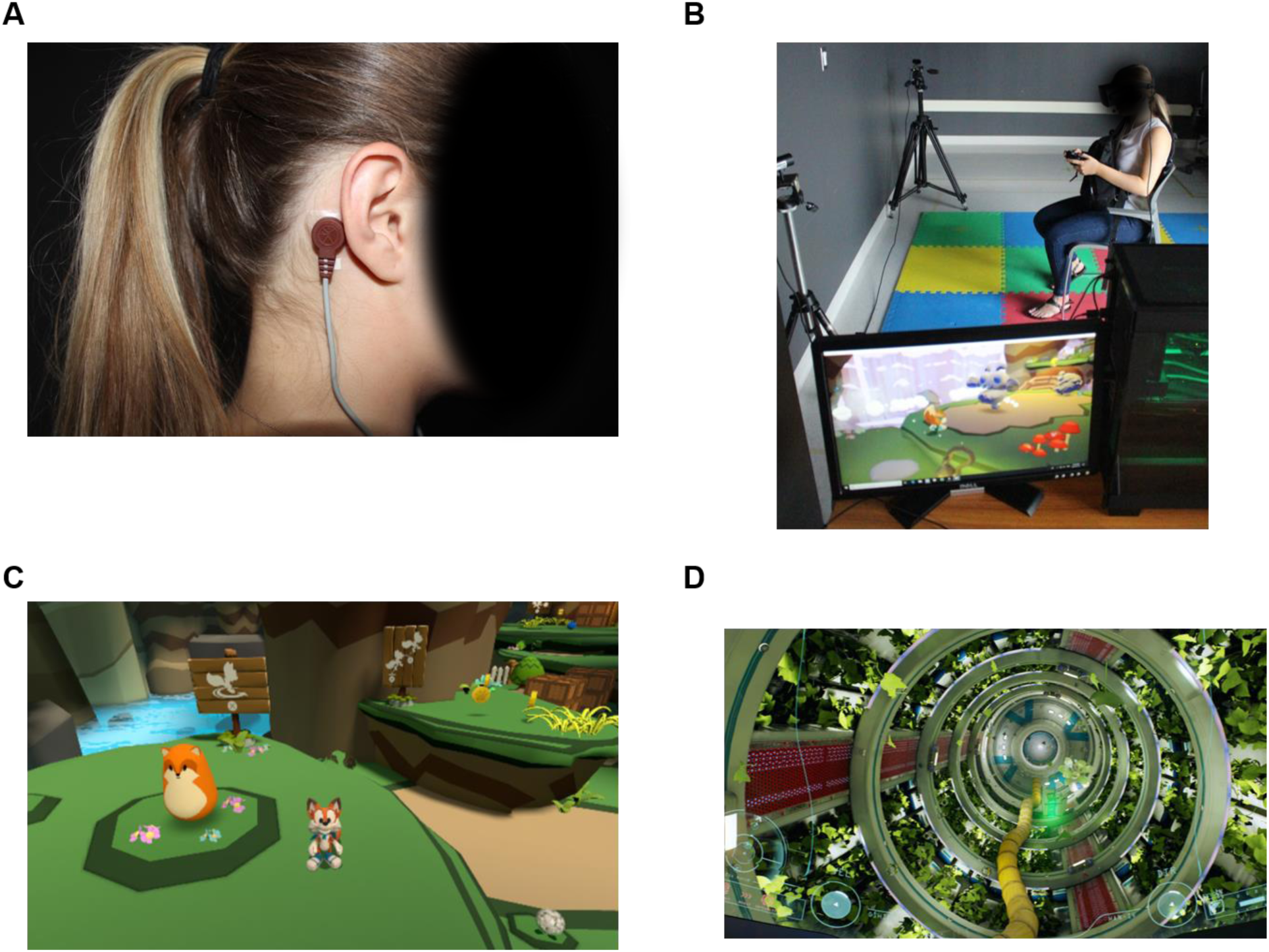
a) GVS electrode affixed at right mastoid process of the participant. b) Illustration of the participant during the VR portion of the study. c) Screenshot from moderate VR content. d) Screenshot from intense VR content

The VR environment was presented with a head mounted display (Rift CV1, Oculus VR; 90 Hz refresh rate, 1080 × 1200 resolution per eye) and the environment was rendered by a high-end graphics card (NVIDIA GTX1070). The headset position was tracked by a combination of inertial (accelerometer/gyroscope) and optical (3 × infrared Oculus cameras) sensors that were part of the commercial device package, and this movement was translated into motion of the observer viewpoint in the VR task. The packaged software of the headset was used to calibrate the capture space (95 × 95 inches) and the inter-pupillary distance of the headset for each participant. A depiction of the setup is shown in Figure 1b.

Participants were exposed to either a ‘moderate’ intensity VR environment (Figure 1c; Lucky’s Tale, version 1.0.2) or an ‘intense’ VR environment (Figure 1d; ADR1FT, version 1.1.8). The two tasks differed with respect to several factors, including the simulated self-motion (the intense task contained greater and more frequent simulated self-motion), visual content (the moderate task contained a cartoon-like aesthetic while the intense task was designed as a photorealistic impression of a space-station), and gravitational reference (no reference for gravity existed in the intense task, where participants were ‘floating’ in zero-gravity, unlike the moderate task). The categorization as ‘moderate’ or ‘intense’ was derived from ratings that are used to advise consumers about VR content on the commercial storefront (Oculus Store). Instructions also differed between the tasks: Participants were asked to advance through the game levels in the moderate task, and were asked to simply explore a space station in the intense task. The participants controlled their movement in either task using a wireless gamepad (XBOX One, Microsoft).

Cybersickness levels were collected using a quick verbal report (Fast Motion Sickness scale, FMS; Keshavaraz & Hecht, 2011) and a questionnaire that was completed after VR exposure (Simulator Sickness Questionnaire, SSQ; Kennedy et. al. 1993). FMS scores are measured as a verbal response to the following question: “On a scale from 0-20 with 0 being no sickness and 20 being severe sickness, how do you feel?” The SSQ is a commonly-used written scale that includes 16 items (e.g., nausea, headache, and eyestrain) that are assigned a severity rating (‘none’, ‘slight’, ‘moderate’, or ‘severe’). Both measures are commonly used in studies of cybersickness and have been validated previously (e.g., Kennedy et. al. 1993; Keshavaraz & Hecht, 2011; Keshavarz et al., 2015a)

### 2.3. Procedure

At the start of the experiment, each participant was shown the GVS device and received verbal instruction on its function. Participants were informed that the effects of GVS differed for each individual, and that the stimulation was imperceptible to some. First the experimenter prepared the skin at the mastoid processes by rubbing briskly with abrasive alcohol gel for approximately 30 sec. Following this, the self-adhesive electrodes were applied to the mastoid processes. Prior to testing, participants were exposed to the GVS stimulus according to their randomly assigned group (GVS or sham; between-subjects factor). The GVS group received a bilateral noisy low-frequency (LF) stimulus at an amplitude of ±1750 µA for a total of 15 s (5 s fade in, 10 s exposure). This form of stimulus consists of a random (normal) current generated at a rate of 1280 samples/s up to and including the designated stimulus level (±1750 µA) which was passed through a digital low-pass filter to significantly dampen any frequencies above 100 Hz. The sham group received a zero-amplitude stimulus for a total of 15 s (5 s fade in, 10 s exposure). The electrode wires were then taped to the participant’s neck to ensure the wires stayed in place during game play. The stimulus amplitude was consistent within each group with no changes made for personal differences. Once the electrodes were attached and the test stimulus was finished, the GVS device was placed in a shoulder backpack on their chest so that the device moved with them. During the period of GVS/sham exposure, the noisy LF stimulus (5 s fade in, peak amplitude of ±1750 µA) was delivered continually over a 30 min period for the GVS group, or zero-amplitude for the sham group. Both groups received brief impedance checks every few seconds that comprised a stimulus amplitude of 120 µA. The maximum permitted impedance level was 55 kΩ, and levels above this value typically indicated that the electrodes were improperly secured.

The procedure was the same between the moderate and intense VR groups. First, participants were asked if they felt well, and if they were currently feeling any sickness prior to starting the VR task. All reported that they felt well and were not experiencing any sickness. Once the participants were ready to enter the designated VR environment, they were shown the controller and all the controls were explained verbally. The participants were seated and underwent the VR headset calibration until visual stimuli were displayed clearly. The study was completed in three phases: first participants completed a 10 min pre-adaptation phase with no GVS stimulus, followed by a 30 min adaptation phase (i.e., the GVS group received noisy vestibular stimulation while the sham group received sham stimulation), and finally a 10 min post-adaptation phase that was identical to the first phase. In each phase, the participant was instructed to engage in the VR task until requested to stop. All phases took place within the same VR environment (either moderate or intense, depending on the group assignment of the participant). During each phase the FMS CS score was taken by the experimenter every 3 min (Keshavarz & Hecht, 2011). After each phase there was a 3 min break during which the SSQ was completed (Kennedy et. al. 1993).

## 3. Results

### 3.1. Demographics and task performance

We examined the effects of participant age, sex, and previous experience with video games/VR on the following sickness outcome measures: participants’ total scores on each SSQ, and participants’ final four FMS scores (i.e., all scores in the post-adaptation block, and the final FMS score from the adaptation block). We found no significant Pearson correlations between age and sickness measures (*p*s ≥ .14), and no difference in sickness scores between men and women (*p*s ≥ .30). In addition, there were no differences between participants who had or had not experienced VR before for any of these sickness measures (*p*s ≥ .09). Although we were unable to analyse the effects of video game experience on sickness scores due to low numbers in each cell, the distribution of individuals who were experienced or inexperienced with video games was relatively equal across groups. For the ‘moderate’ VR task, we ran Pearson correlations to assess the association between the number of levels of the VR game participants completed and the same sickness measures as above, and found no significant relationships (*p*s ≥ .07).

### 3.2. Baseline effects

We first determined if there were any baseline differences between the GVS and sham stimulation groups by conducting a mixed design analysis of variance (ANOVA) on FMS scores for the between-subjects factor of stimulation (GVS or sham) and the within-subjects factor of FMS repetition (1, 2, or 3) in the pre-adaptation block. We observed no effects of stimulation group, and no interaction between the factors for the intense VR content group (*p*s ≥ .10) or the moderate VR content group (*p*s ≥ .12). For the moderate VR content group, we observed no effect of FMS repetition in the pre-adaptation block (*p* = .12), whereas there was an increase in FMS scores over time for the intense VR content group (*F*(2, 36) = 7.05, *p* = .003, *GES* = .08).

### 3.3. Adaptation effects

We computed the effect of exposure to stimulation as the post-adaptation FMS scores minus the pre-adaptation FMS scores. We subtracted these scores item-wise, that is, the first FMS score in the pre-adaptation block was subtracted from the first score in the post-adaptation block, and so on. For this dependant variable (‘adaptation effect’), we conducted mixed design ANOVAs (between-subjects factor, stimulation: GVS or sham; within-subjects factor, FMS repetition: three levels).

#### Intense VR content

We found a significant main effect of stimulation (*F*(1, 18) = 4.47, *p* = .048, Generalized Eta Squared (GES) = .14), but no effect of FMS repetition (*F*(2, 36) = 0.41, *p* = .66, GES = .007), and no interaction between the two factors (*F*(2, 364) = 120, *p* = .31, GES = .02). Figure 2a depicts these data.

**Fig. 2.**
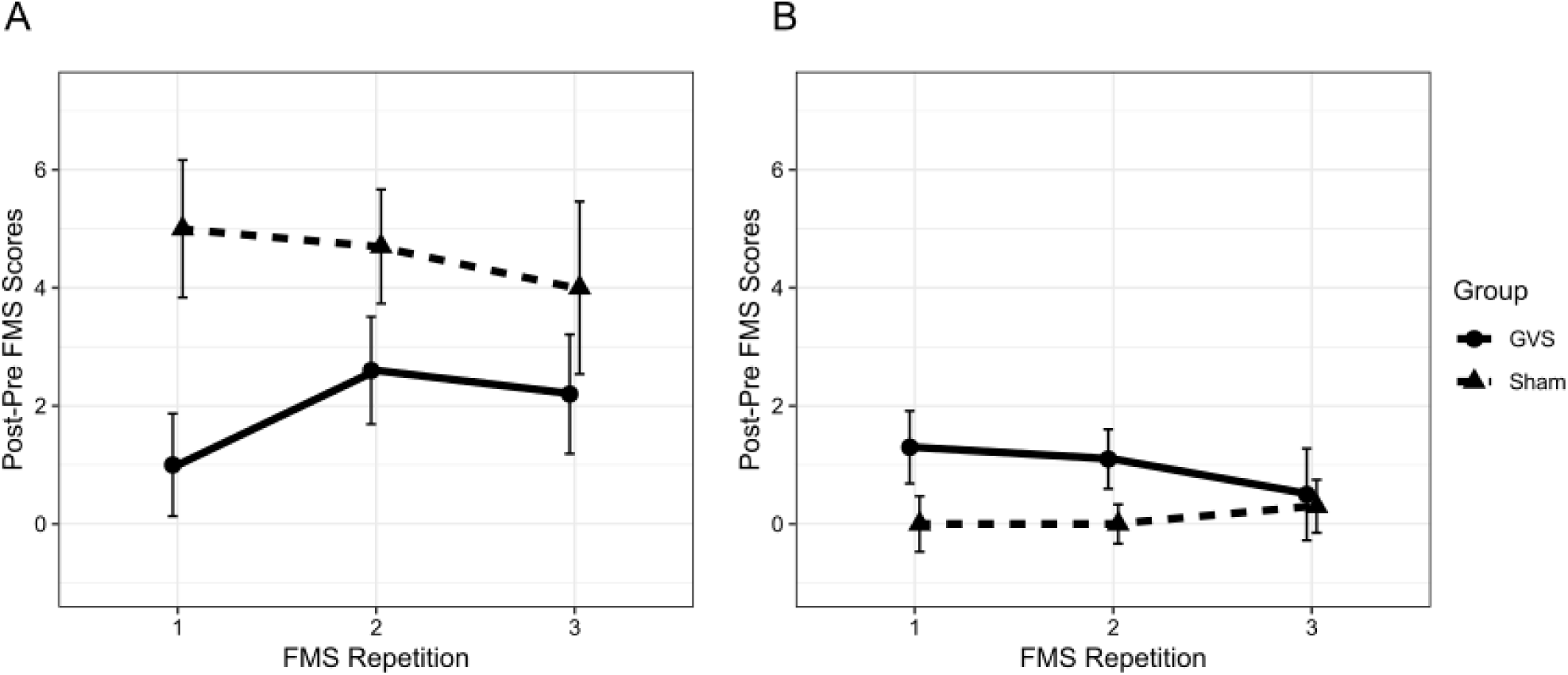
Adaptation effects (plotted as post minus pre adaptation FMS scores) over FMS repetitions for groups who experienced (a) the intense VR content, and (b) the moderate VR content. Error bars are standard errors

We conducted a follow-up analysis consisting of least-squares means *t*-tests with a conservative Satterthwaite adjustment to the degrees of freedom (Lenth 2016; note that non-parametric tests revealed similar results). Examining the effect of stimulation as a function of FMS repetition revealed a significant difference between the GVS and sham groups at the first FMS repetition (*t*(36.2) = 2.61, *p* = .01, Cohen’s *d* = 1.31), but not at the second or third FMS repetitions (*p*s ≥ .18).

Polynomial analysis showed no significant trends for the adaptation scores across FMS repetitions in the GVS or sham groups (*p*s ≥ .25).

#### Moderate VR content

The same mixed-design ANOVA on post-pre FMS scores revealed no significant differences between GVS and sham groups, no effect of FMS repetition, and no interaction effect (*p*s ≥ .18). Figure 2b depicts these data.

### 3.4. Effects during GVS exposure

#### Intense VR content

Inspecting the FMS scores from the adaptation block alone (mixed design ANOVA, between-subjects factor, stimulation: GVS or sham; within-subjects factor, FMS repetition: ten levels) revealed that there was no main effect of stimulation (*F*(1, 18) = 1.12, *p* = .30, GES = .04), but we observed a significant effect of FMS repetition (*F*(9, 162) = 7.33, *p* < .001, *GES* = .11), and an interaction between the two factors (*F*(9, 162) = 3.54, *p* < .001, *GES* = .06).

Follow-up analysis of the interaction effect using least squares means *t*-tests with Satterthwaite-adjusted *df* (Lenth 2016) revealed that the GVS group had significantly lower FMS scores than the sham group only at FMS repetition 10 (*t*(36.8) = 2.98, *p* = .005, Cohen’s *d* = 1.49; all other *p*s ≥ .07). Figure 3a depicts these data.

**Fig. 3.**
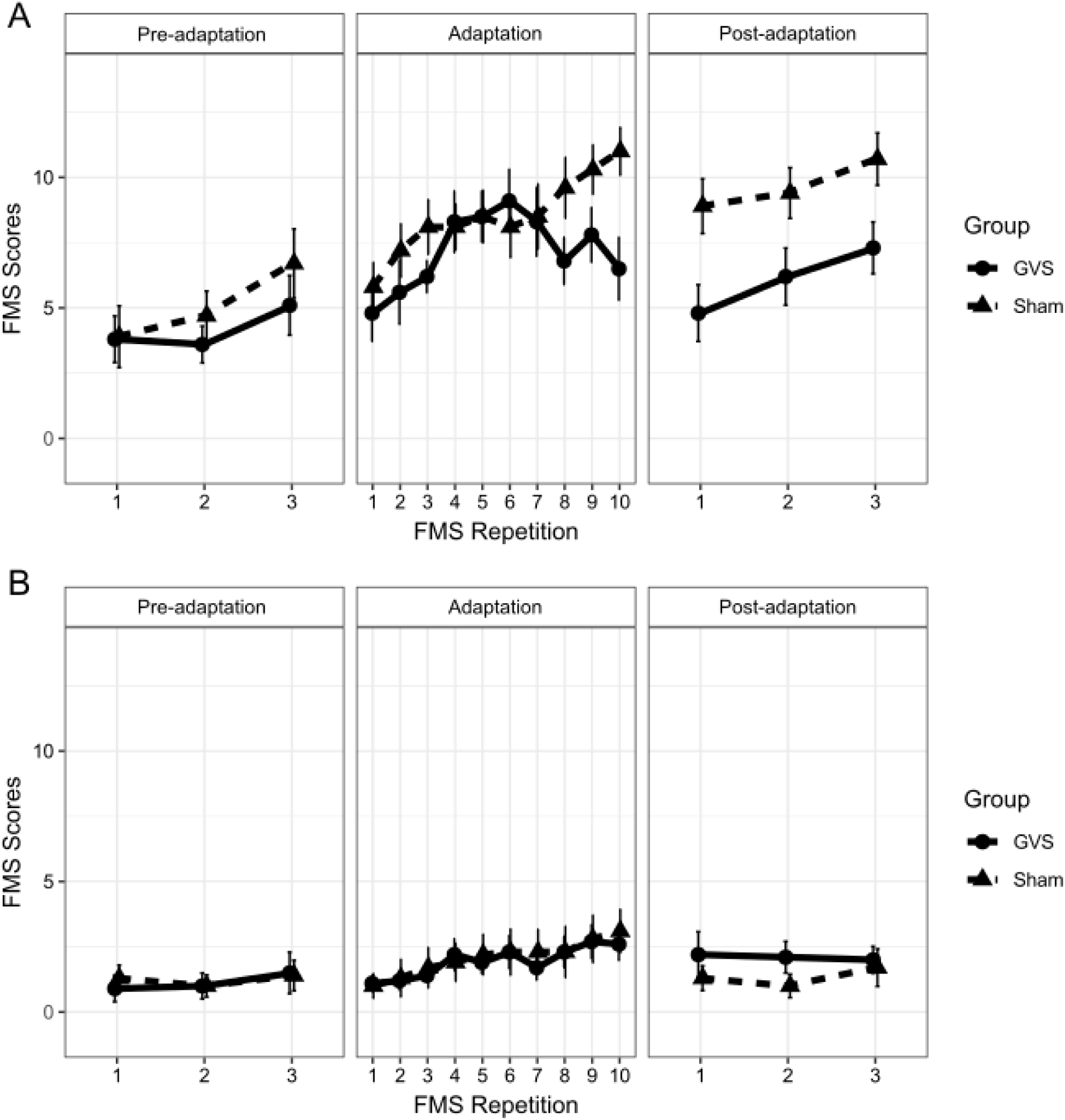
FMS scores by stimulation group across repetitions in the three phases of the experiment for groups who experienced (a) the intense VR content, and (b) the moderate VR content. Error bars are standard errors

We also observed significant linear trends for both the GVS group (*t*(81) = 2.55, *p* = .01) and sham group (*t*(81) = 8.73, *p* < .001) during the adaptation block. Importantly, FMS scores for the GVS group also tended to decrease during the later parts of the block, reflected in a significant quadratic trend for the GVS group (*t*(81) = 4.54, *p* < .001) that did not emerge for the sham group (*t*(81) = 0.34, *p* = .73).

#### Moderate VR content

A mixed-design ANOVA revealed that FMS scores from the adaptation block differed across FMS repetitions for the moderate VR content group (*F*(9, 162) = 6.95, *p* < .001, *GES* = .07), but there was no main effect of stimulation, and no interaction effect (*p*s ≥ .86). The tendency for FMS scores to increase across FMS repetitions was borne out in significant linear trends for both the GVS group (*t*(81) = 4.55, *p* < .001) and the sham group (*t*(81) = 6.16, *p* < .001). No higher-order trends were observed for either group (*p*s ≥.56). Figure 3b depicts these data.

### 3.5. Simulator sickness questionnaire

The results of the SSQ indicated that there was no difference between the GVS and sham groups following the intense VR content. This was the case for all three completions of the SSQ, both for the moderate and intense VR content groups (*p*s > .05).

### 3.6. Symptomatology

To assess the symptomatology of cybersickness, we computed the total number of participants that reported each symptom on the SSQ (any severity) from the ‘adaptation’ and ‘post-adaptation’ blocks and identified the most common symptom (or symptoms) for each condition and group. The results are depicted in Table 1. The pattern of symptoms experienced in each condition was similar, with eyestrain, general discomfort, and fatigue being the most common symptoms across participants.

**Table 1.**
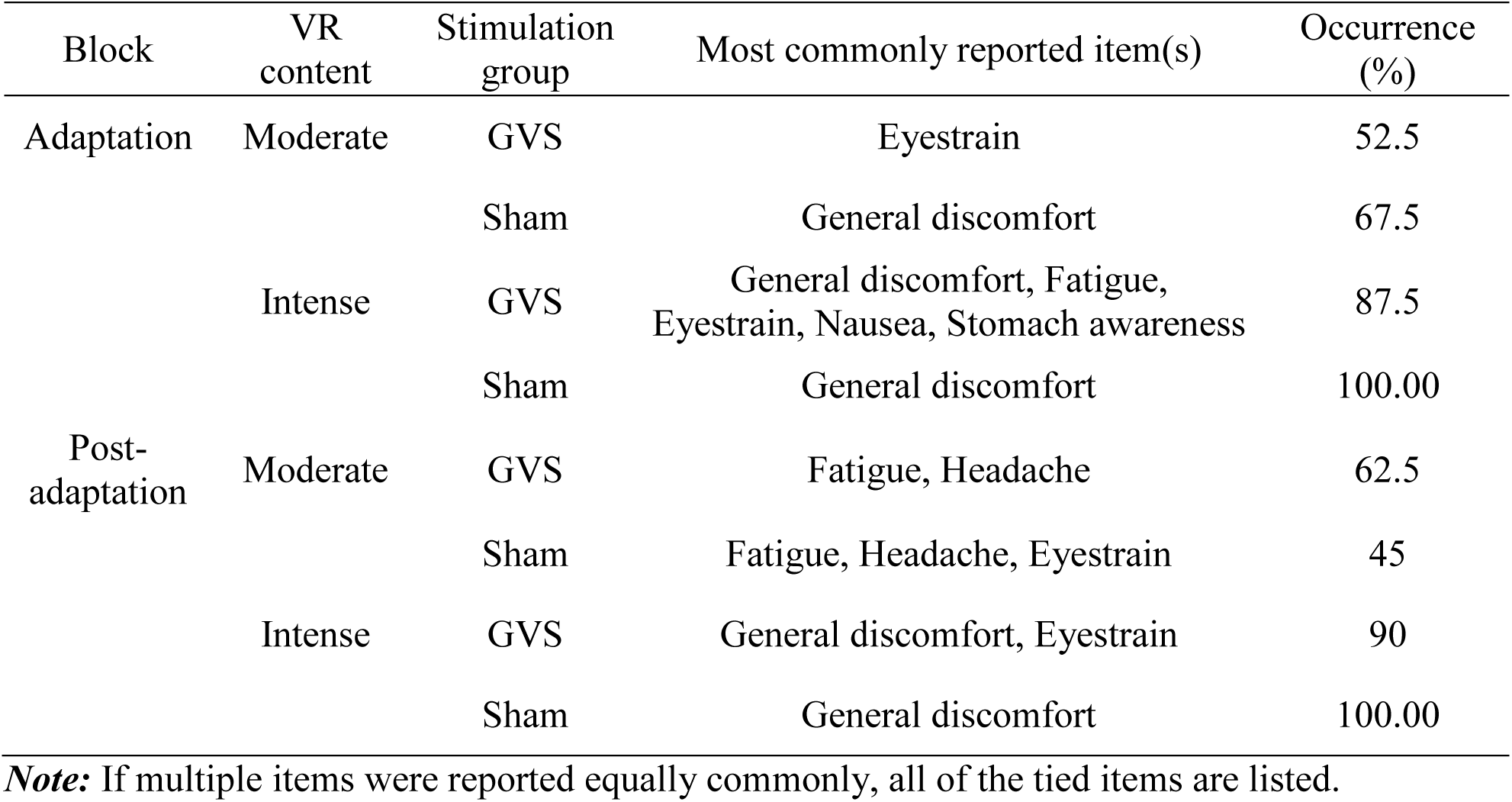
Symptomatology Derived from the SSQ across Groups and Conditions.

| Block | VR content | Stimulation group | Most commonly reported item(s) | Occurrence (%) |
| --- | --- | --- | --- | --- |
| Adaptation | Moderate | GVS | Eyestrain | 52.5 |
|  |  | Sham | General discomfort | 67.5 |
|  | Intense | GVS | General discomfort, Fatigue, Eyestrain, Nausea, Stomach awareness | 87.5 |
|  |  | Sham | General discomfort | 100.00 |
| Post-adaptation | Moderate | GVS | Fatigue, Headache | 62.5 |
|  |  | Sham | Fatigue, Headache, Eyestrain | 45 |
|  | Intense | GVS | General discomfort, Eyestrain | 90 |
|  |  | Sham | General discomfort | 100.00 |

**Note:** If multiple items were reported equally commonly, all of the tied items are listed.

## 4. Discussion

Here we show evidence that the addition of vestibular noise during VR exposure is associated with a reduced severity of CS. Given previous evidence that CS is reduced by bone-conducted vibration at the mastoids (Weech et al., 2018a), we expected that noisy galvanic stimulation would also reduce CS. One possible mechanism for this effect involves the down-weighting of ‘unreliable’ vestibular cues by the nervous system, thus up-weighting vision and reducing the impact of sensory conflicts in VR, consistent with optimal cue integration theory (Ernst & Banks, 2002; Weech & Troje, 2017; Weech et al., 2018b). While the SSQ scale revealed no differences between the stimulation groups, pre-minus-post FMS scores were significantly lower for the GVS group compared to the sham group. The reduction in CS was very transient (3-6 min), providing preliminary evidence that noisy vestibular stimulation may be a viable method for improving user comfort in VR. At the same time, no effect of GVS was observed during a mildly provocative VR application, confirming that there are minimal effects on comfort when noisy GVS is used during a comfortable experience. An informal comparison of symptomatology across stimulation groups also suggests noisy GVS is unlikely to generate major discomfort.

One possible explanation of our results involves recalibration of visual and vestibular weights at the level of the cerebellum due to noise exposure (Dilda et al., 2014; Ernst & Bülthoff, 2004; Moore et al., 2015). The absence of GVS-related effects for the moderate VR content suggests that the intervention might be effective only when frequent sensory conflicts related to self-motion occur, although on this point we are limited to speculation given that there were several other differences between the moderate and intense tasks (e.g., task instructions, visual style, the presence of a gravitational reference). While there are several aspects of the observed effects that are currently unclear, the results provide further support for the utility of noisy stimulation for reducing the severity of CS.

### 4.1. Practical significance

The magnitude of the benefit provided by noisy GVS is conveyed by the large effect size (Cohen’s *d* = 1.3) for the difference between the GVS and sham groups immediately following the adaptation block for the intense VR content. For the GVS group, the difference between post- and pre-adaptation FMS scores was approximately zero at the first FMS repetition, suggesting that the accumulated effects of nauseogenic VR exposure (demonstrated by significant linear trends in other conditions) were practically eliminated for these participants. The benefit of GVS was also much larger here than in a previous study where noisy vestibular cues were applied using another technique (bone conducted vibration) and where the authors identified only a ‘medium’ effect size (η^2^_*p*_ = 0.20; Weech et al., 2018a).

However, the practical significance of these results is limited in some regards. For instance, given that the effect of GVS had washed out after an additional 3 minutes of VR exposure, there may be a limited benefit of the current iteration to users in a practical setting. Other research has identified long-lasting effects (> 6 months) of multi-session noisy GVS on postural control and perceptual upright estimation (Dilda et al., 2014; Moore et al., 2015). Both of these tasks require the integration of multisensory estimates of the state of the world and the body, and the sustained effects of noisy GVS observed in these studies suggest that noisy stimulation caused down-weighting of vestibular cues that would otherwise interfere with accurate completion of the task. Therefore, it is conceivable that CS–which is often attributed to a problem with disregarding inaccurate vestibular cues (e.g., Weech et al., 2018a)–may also be mitigated by long-term exposure to noisy stimulation (i.e., several sessions over multiple days). This proposition remains to be tested in future research. It also remains to be seen if visual noise manipulations result in a symmetric effect on CS, such that visual noise delays the process of reaching visual dominance for self-motion perception, resulting in more sensory conflicts. While evidence suggests that visual brightness and contrast manipulations do not affect CS (Shahal et al., 2016), other manipulations should be tested (e.g., visual motion coherence, blurring) before conclusive statements can be made about visual noise and CS.

The magnitude of current applied in the GVS group was consistent across participants. Although there was large variability across participants in this group with respect to the reduction in CS, the significant effects we observed suggest that minimal individualisation of the stimulation parameters may be required in order to achieve a benefit. It is not clear how much greater the reduction in CS might have been if individual variability in vestibular sensitivity (which can vary considerably; e.g., Grabherr et al., 2008; Soyka et al., 2012, 2011) were taken into account in the range of current values that were applied during stimulation. Future efforts to modulate the magnitude of vestibular noise as a function of vestibular thresholds may prove useful in reducing the inter-individual variability we observed.

There have been several attempts to mitigate the negative effects of VR use (e.g., Cevette et al., 2012; Dorado & Figueroa, 2014; Fernandes & Feiner, 2016; Gálvez-García et al., 2015; Kim et al., 2008; Reed-Jones et al., 2007; Stanney & Hash, 1998; Klüver et al., 2015), but the practicality of current interventions is limited (Weech et al., 2018a). One constraint of previous approaches to reducing CS based on visual-vestibular recoupling has been the need to couple vestibular stimulation to the visual self-motion cues depicted by the simulated environment (e.g., Cevette et al., 2012; Reed-Jones et al., 2007; Weech et al., 2018a). The need to couple the stimulation device to the rendering software results in computational expense, and also results in potentially important delays between the initial generation of the appropriate stimulus on the rendering hardware, and the stimulus reaching the stimulation device at the user. The noisy GVS approach is therefore appealing in a practical sense, as it requires no software-coupling in order to achieve the significant reduction in CS that we observed here.

### 4.2. Discrepancy between cybersickness measures

While the FMS scale revealed differences between the stimulation groups, the SSQ results did not differ between the two groups. The reason for this discrepancy between measurements is not clear, especially since the effects of GVS persisted between the adaptation and post-adaptation phases—during which one of the SSQ scales was completed. It is possible that a replication of this study that uses a larger sample size would help to reveal similar effects on SSQ scores. One open question concerns the extent to which probing participants for CS reports during a VR task (FMS scores), which we did repeatedly here, may affect the either physiological experience of CS or give rise to a response bias that depends on previous responses. Upcoming experiments should address these issues by examining physiological and subjective reports both with and without intermittent prompting.

### 4.3. Other theories of cyber/motion sickness etiology

We have discussed the prospect that vestibular noise reduced CS by mitigating sensory conflict, although it should be noted that the results could equally be aligned with other etiological accounts of CS. In line with the postural instability account of motion sickness (Riccio & Stoffregen, 1991; Stoffregen et al., 2014; Smart Jr. et al., 2002), which links the use of maladaptive control strategies to sickness etiology, it is conceivable that noisy GVS influenced postural control strategies, resulting in more adaptive control of the head and body for the GVS group than for the sham group. In order to test this proposition, it would be useful to run further examinations of the head movements of the GVS and sham group participants, although these data were not collected here. However, it is clear that there are few tests that could possibly distinguish between the postural instability/sensory re-weighting accounts of CS (Nishiike et al., 2013; Weech et al., 2018a). The ability to integrate multiple sensory cues is a key aspect of stable bodily control, and this inextricable link between postural control and sensory dynamics means that any measurement of one factor is likely to co-vary with measurements of the other. Future studies that attempt to partition the effects of postural control and sensory re-weighting will likely prove crucial to the refinement of CS reduction approaches.

The use of GVS in the current study could also have resulted in changes in sensory processing via stochastic resonance (SR), where adding noise to a sensory channel elevates the detection of true signals (Moss et al., 2004; Wiesenfeld & Moss, 1995). Others have shown increased postural stability resulting from low-amplitude noisy current applied to the vestibular system, perhaps due to facilitated detection of head movement (Mulavara et al., 2011; Pal et al., 2009). However, the amplitude of current used in the current study (±1750 µA) is higher than that used in these previous studies (amplitude is commonly in the range of ±100 to ±500 µA), and low-amplitude current is a key requirement for SR-like effects to emerge. In addition, SR effects typically emerge rapidly, and do not demonstrate an adaptation period such as the one we observed here. As such, we do not expect that SR played a significant role in the current study.

In a similar manner, it is conceivable that noisy GVS reduced cybersickness by increasing the level of presence experienced by participants in the VR environment. Presence and CS share an inverse correlation, whereby high presence seems to impart a protective effect on VR users (Weech et al., 2019). This explanation of our results would also be consistent with empirical evidence showing that vection is facilitated by noisy GVS (Weech & Troje, 2017), given the strong relationship between vection and presence in VR (Keshavarz et al., 2015b; Weech et al., 2019). Since we did not measure presence in the current study, we cannot identify whether the effects observed can be explained by this related factor. A future replication should address this prospect. Other extraneous factors could also be measured in a replication, including distraction (did GVS distract participants, thus reducing their awareness of CS?),

It is also possible that electrical stimulation of the vestibular apparatus produced changes in the velocity storage process. As documented by Cohen and colleagues (2003, 2008; Dai et al., 2007, 2010, 2011), motion sickness susceptibility is associated with processing differences in the vestibular pathway that stores information about low-frequency head motion and the gravitational vertical (i.e., vestibular velocity storage). Repeated passive movement of the head (Cohen et al., 1992) and electrical stimulation of the vestibular cerebellum (Solomon et al., 1994) or 5 mA bipolar GVS (Karlberg et al., 2000) are sufficient to change the time constant of velocity storage, likely by stimulating GABAergic inhibition of cells in the vestibular nuclei that are involved in storing velocity estimates. Accordingly, it is conceivable that the effects observed here might be attributable to changes in velocity storage following galvanic stimulation. While the effect of noisy GVS on the velocity storage mechanism is currently unexplored, this explanation of the current findings cannot be ruled out.

## 5. Conclusion

Here we report evidence that noisy galvanic vestibular stimulation applied during an intense VR application can result in a transient reduction in CS severity. The side-effects of GVS on CS symptomatology were minimal and did not increase discomfort in a non-intense VR application. The washout of the effects was very rapid, indicating a fast re-establishment of the un-adapted state of vestibular processing. We posit that the effects observed here might be extended upon longer durations of exposure to noisy GVS, similar to the long-lasting effects observed for GVS-induced sensory re-weighting and postural adaptation (Dilda et al., 2014; Moore et al., 2015). Overall, the findings demonstrate the potential for CS reduction by way of a non-invasive sensory stimulation method that requires no software-coupling and minimal efforts to achieve personalisation, although further refinements of the method used here will be required in order to establish its practical utility.

## Acknowledgements

This research was supported by grants to MBC from Oculus Research, the Ontario Research Fund and Canadian Foundation for Innovation’s John R. Evans Leaders Fund, and the Natural Sciences and Engineering Research Council of Canada. The industry sponsor had no influence in the design or execution of the current research. All authors declare that there are no conflicts of interests.

